# A superfamily 1 helicase translocates on double-stranded DNA

**DOI:** 10.64898/2026.09.16.752024

**Authors:** Sushil Pangeni, Amatullah Mustafa Nakara, Olivia Yang, Fahad Rashid, James M. Berger, Taekjip Ha

**Affiliations:** Program in Cellular and Molecular Medicine, Boston Children’s Hospital, Boston, Massachusetts, USA; T. C. Jenkins Department of Biophysics, Johns Hopkins University, Baltimore, Maryland, USA; Department of Biophysics and Biophysical Chemistry, Johns Hopkins University School of Medicine, Baltimore, Maryland, USA; Department of Pediatrics, Harvard Medical School, Boston, Massachusetts, USA; Howard Hughes Medical Institute, Boston, Massachusetts, USA

## Abstract

Replicative and accessory helicases are essential for maintaining genome integrity during DNA replication and repair. Rep helicase belongs to superfamily 1 (SF1), the first helicase family solved at atomic resolution and a longstanding model for how helicases couple ATP hydrolysis to movement on DNA. Using optical tweezers combined with fluorescence imaging, we characterize how Rep translocates and navigates diverse nucleic acid and protein obstacles. We directly observe highly processive translocation on ssDNA (∼300 nt/s over >13,000 nt) and show that this directional movement can drive single-stranded DNA-binding proteins (bacterial SSB, archaeal SSB, and eukaryotic RPA) along ssDNA without slowing. On encountering a short duplex barrier (≤60 bp), either a DNA duplex or an RNA/DNA hybrid, Rep frequently bypasses it, with termination and unwinding also observed. Unexpectedly, we identify a mode in which Rep transitions from ssDNA onto dsDNA and translocates directionally along the duplex with high processivity(>12,000bp) and at a much higher speed than on ssDNA (∼1100 bp/s). AlphaFold3 modeling is consistent with Rep tracking a single strand while keeping the duplex intact. To our knowledge, this is the first demonstration of processive dsDNA translocation by an SF1 helicase, and it broadens models for how translocases operate on DNA during replication and repair.

## Introduction

Helicases are ATP dependent motor proteins that unwind duplex nucleic acids and resolve DNA and RNA secondary structures^1^. Decades of biochemical, structural, and single molecule studies have revealed extensive mechanistic details for many helicases involved in DNA and RNA metabolism^2–8^. Based on conserved structural motifs and directional movement along nucleic acid substrates, translocases are defined as helicases whose core function is not replicative fork unwinding but includes ssDNA translocation, DNA unwinding, protein displacement, and other activities that support replication and repair^7^. In *E. coli*, the replicative helicase DnaB is a hexameric ring motor essential for fork progression, whereas other Superfamily 1 (SF1) helicases, specifically the 3’ to 5’ translocating SF1A subgroup that includes Rep, UvrD, and DinG (SF1B enzymes move 5’ to 3’)^9^, function as translocases^2,10^.

Rep helicase is among the most extensively characterized molecular motors in *E. coli*, with roles spanning DNA repair, replication, and replication restart. Its conformational cycle, subdomain functions, and ATPase coupled stepping mechanism have been dissected through structural studies, mutagenesis, and chemical crosslinking^1,8,11^. In solution, Rep is monomeric and unwinds duplex DNA only weakly, often to an undetectable degree; efficient unwinding requires activation, achieved by a partner protein, by crosslinking that stabilizes the unwinding-active closed conformation^11^, or by dimerization^12^, which recent structural studies indicate acts by repositioning the regulatory 2B subdomain^13^. Early measurements of its single stranded DNA (ssDNA) translocation speed and processivity were performed using stopped flow assays on substrates of up to 124 nt and were later corroborated by single molecule FRET experiments on surface tethered DNA^14,15^. Despite this wealth of biochemical and structural information, how Rep operates within the crowded, protein rich environment likely to be found in a replication fork or repair intermediate remains unclear.

Movement along double-stranded DNA is itself a known motor activity, but one associated with other enzymes: many SF2 members, including the chromatin and fork-remodeling enzymes Rad54, SMARCAL1, and HLTF, translocate along intact duplex DNA without unwinding it^16^, as do ring-shaped helicases. SF1 helicases, by contrast, have been regarded strictly as ssDNA translocases. In the cell, Rep contributes to replication and to the restart of stalled replication forks, and helps replication proceed through regions where it collides with transcription or with tightly bound proteins^17–20^. Yet how a translocase navigates dsDNA, DNA–RNA hybrids, protein roadblocks, or structured DNA, and whether and how Rep transitions between ssDNA and dsDNA during these events, has not been directly observed. Observing these encounters in real time, one by one in vitro, strengthens the mechanistic understanding of how these complexes may behave in cells.

Single-molecule methods have been essential for dissecting how DNA motors move, because they resolve pauses, backtracking, heterogeneity, and rare events that ensemble measurements average away. Rep itself illustrates this: single-molecule fluorescence first showed that a Rep monomer translocates 3^′^ to 5^′^ on ssDNA and repetitively shuttles upon meeting a blockade^15^, and cross-linking a Rep monomer into its unwinding-active conformation converts it into a processive superhelicase^11^. More recently, instruments that combine optical trapping with fluorescence imaging have allowed motor proteins to be followed directly on long DNA under defined tension; this approach revealed, for example, that the eukaryotic replicative helicase CMG can transition between single- and double-stranded DNA at a fork^21^.

Here, we use a combined optical trap and multicolor confocal fluorescence microscope to monitor Rep translocation on kilobase length ssDNA in real time while simultaneously probing its interactions with defined nucleic acid and protein obstacles. This system enables controlled reconstruction of the diverse challenges a translocase may encounter, including short dsDNA segments, replication fork-like structures, and bound proteins. Contrary to the expectation that SF1 helicases function exclusively as ssDNA translocases that unwind, rather than traveling along, any duplex they encounter, we find that Rep can transition onto dsDNA and translocate directionally along duplex substrates over thousands of base pairs. Structural modeling using AlphaFold3 suggests an inchworm mechanism analogous to its ssDNA stepping cycle but on an intact duplex. Together, these observations provide new insight into how Rep and potentially other bacterial translocases navigate complex nucleic acid landscapes during their function.

## Results

### Direct observation of Rep translocation on ssDNA

We first established a quantitative baseline for Rep movement on ssDNA. Rep contains the canonical SF1A subdomains 1A, 1B, 2A, and 2B (Figure 1B)^22^. Purified Rep unwound a 30-bp duplex containing a 3^′^ overhang in an ensemble FRET assay, as indicated by the increase in donor fluorescence upon ATP addition to a solution containing Rep and DNA (Figure 1C). For single-molecule imaging, we used the previously described double-cysteine Rep (Cys178/Cys400) labeled with a mixture of Cy3 and Cy5^15^; a signal in either fluorescence channel was used to identify binding and movement events.

**Figure 1:**
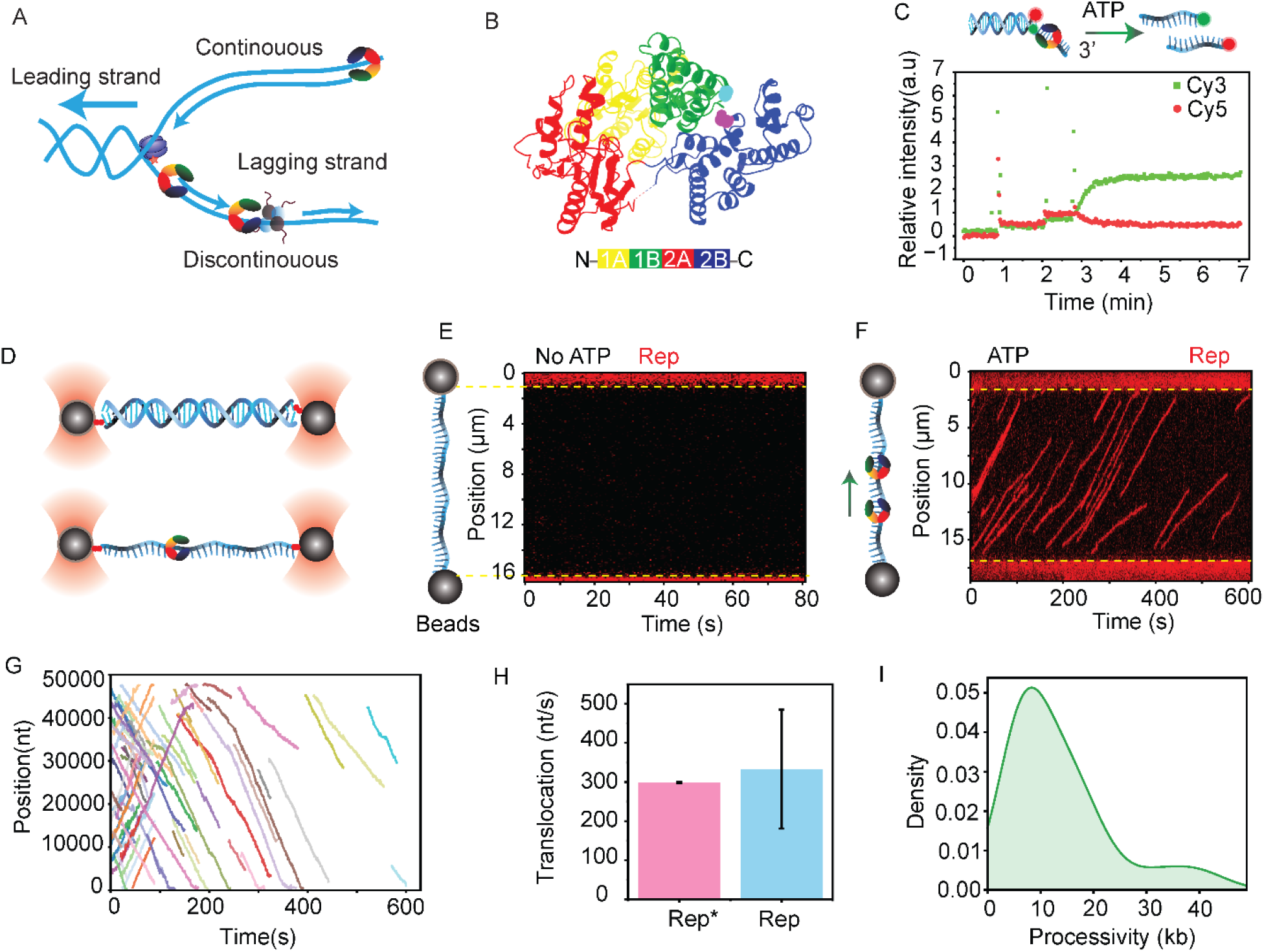
Rep is a ssDNA translocase. **A)** Schematic of a replication fork. The leading strand is continuously synthesized, driven by unwinding by the replicative helicase DnaB. The lagging strand is synthesized discontinuously, leaving ssDNA platforms that bind SSB and accessory translocases like Rep. **B)** Rep helicase structure (PDB: 1UAA) with the established 1A, 2A, 1B, and 2B domains labeled accordingly. **C)** Ensemble FRET assay establishing Rep helicase activity. **D)** Optical trap schematic used to study Rep translocation. Top: dsDNA tethered to streptavidin-coated beads via terminal biotins. Bottom: ssDNA generated by overstretching dsDNA as shown in the top panel. Rep binds to ssDNA and its translocation is observed by confocal scanning. **E)** Representative kymograph of Rep in the absence of ATP and Mg^2+^. **F)** Representative translocation kymograph of Rep on ssDNA over time in the presence of ATP and Mg^2+^. **G)** Combined trace plot for Rep translocation at one condition (1 mM ATP, N = 69). **H)** Quantification of Rep translocation rate (N = 69, error=SDV). Rep DM4 represented as Rep is the variant used in this study. Translocation rate is compared to the published Rep translocation rate reported in Myong et al., Science 310, 1144–1147 (2005) represented as Rep*. I)Processivity of Rep helicase on ssDNA as depicted for the traces in G (N=69).

To visualize Rep movement in real time, we used a dual-trap optical tweezers instrument coupled to multicolor confocal fluorescence detection (LUMICKS C-Trap)^21,23-26^. A lambda phage DNA (λDNA) molecule was tethered between streptavidin-coated beads through biotins attached to the same strand. Overstretching removed the complementary strand and generated an ssDNA tether^27^ (Figure 1D), which was subsequently held at 5 pN unless otherwise stated. In the absence of ATP and Mg^2+^, labeled Rep did not bind detectably under our imaging conditions (Figure 1E). With ATP and Mg^2+^, Rep loaded onto ssDNA and showed sustained, directional movements along the ssDNA tether with occasional pauses and restarts (Figure 1F–G). Although our assay does not directly determine strand polarity, the well-established 3^′^ to 5^′^ directionality of Rep and related SF1A translocases^15^ allows us to infer that the observed movement corresponds to 3^′^→ 5^′^ translocation. The mean rate of ssDNA translocation was ∼300 nt/s (n=69) (Figure 1H), similar to reported values^14-15^. The observed trajectories had a mean travel length of 10,200 nt (n=69) (Figure 1I). These measurements establish the rate and travel-length distributions used for comparison with the barrier assays below.

### Rep drives the directional movement of ssDNA-binding proteins

We next examined the movement of ssDNA-binding proteins when they are encountered by a ssDNA-translocating Rep. We used a five-channel assay to load a fluorescent ssDNA-binding protein onto ssDNA and then transfer the tether to a separate channel containing unlabeled Rep, allowing us to visualize the outcome of such an encounter (Figure 2A-2B).

**Figure 2:**
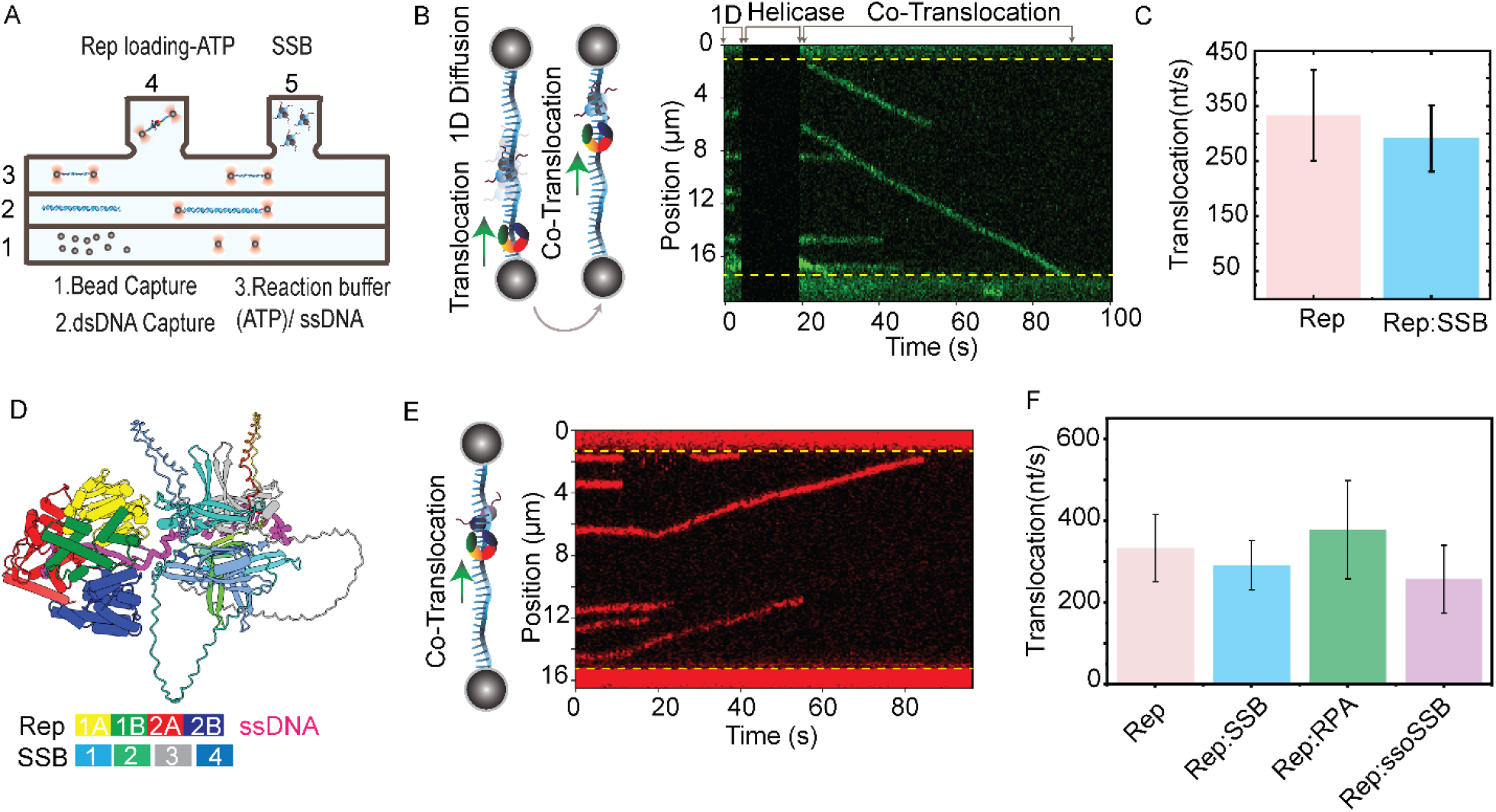
Rep interacts with other accessory proteins. **A)** Optical trap five-channel configuration used to monitor interactions between Rep and SSB-like proteins. **B)** Left: Schematic illustrating how ssDNA-binding proteins diffuse in 1D along ssDNA and how they may encounter a directionally translocating Rep helicase. Right:Representative kymograph showing EcoSSB being displaced by Rep on ssDNA. **C)** Quantification of translocation rates during co-translocation with EcoSSB (N = 16) or with Rep alone. **D)** AlphaFold3 prediction of Rep helicase and SSB bound to a 33-nt ssDNA substrate. **E)** Representative kymograph of Rep pushing SsoSSB along ssDNA. **F)** Comparison of Rep co-translocation with SSBs across species: EcoSSB (N = 16), SsoSSB (N = 17), and RPA (N = 32).

We first tested *E. coli* single stranded DNA binding protein (ecoSSB), which undergoes one ⍰dimensional diffusion on ssDNA^28^. We used labeled SSB^AF555^ (a cystine mutant, A122C SSB(A122C), was labeled using Alexa Fluor 555 maleimide)^29^ and unlabeled Rep. Since SSB undergoes random 1D diffusion^28–32^ and Rep translocates directionally (Figure 1), directional movement of SSB would be the result of helicase pushing (or pulling). Because Rep was unlabeled, only SSB was directly visualized; upon moving the construct with SSB into the channel containing Rep and ATP, a subset of SSBs transitioned from random diffusion to directional, processive translocation. We interpret these events as Rep-driven directional SSB movement (Figure 2B). When multiple SSB molecules underwent processive movements on the same DNA molecule, they moved in the same direction, further supporting the interpretation of co-translocation. Such co⍰ translocation frequently persisted for tens of kilobases without an obvious pause. Translocation rates showed no clear differences between Rep alone and ecoSSB movements induced by Rep (Figure 2C), indicating that SSB does not impose a detectable kinetic penalty for Rep translocation under our conditions.

To ask whether simultaneous binding was sterically plausible, we generated an AlphaFold3 model of a Rep–SSB–ssDNA complex (Figure 2D). This model suggested that both proteins can simultaneously engage the ssDNA, with Rep contacting the backbone and SSB wrapping the ssDNA in its characteristic manner. Although a limited set of Rep-SSB contacts was observed at the interface (e.g., involving 1A subdomain residues Arg209 and Glu199), we did not find evidence for an extensive, high-confidence protein-protein interface, consistent with a scenario in which the directional movement of Rep rectifies the spontaneous diffusion of SSB^29,31,33-34^ rather than relying on a strong protein–protein interaction.

We next tested SSBs from other organisms. Despite differences in oligomeric state and sequence, SSBs share similar OB ⍰fold architecture ^35-36^ and several SSBs have been shown to undergo diffusion on ssDNA^28,30,37^. Using SSB from an archaeal organism, *Sulfolobus solfataricus* (ssoSSB ^Atto650^), we observed transition from diffusional to directional movement of ssoSSB induced by Rep (Figure 2E), again at a rate with no measurable difference from Rep’s translocation rate (Figure 2F). Finally, we tested yeast RPA ^MB543^, a eukaryotic ssDNA⍰binding protein with distinct domain organization and binding modes. RPA also moved directionally in the presence of Rep and ATP at a rate that is comparable to Rep’s translocation rate (Figure 2F).

Together, these results show that Rep can drive the directional movement of a variety of ssDNA binding proteins including bacterial SSB, archaeal SSB, and eukaryotic RPA along ssDNA without a detectable reduction in translocation speed. Given that these proteins undergo diffusion on ssDNA, and that AF3 models do not predict an extensive Rep–SSB interface, the most likely explanation is that Rep’s directional motion biases the diffusive movement of the bound protein. This rectification mechanism is consistent with prior observations of long range co translocation between human RPA and the structurally homologous yeast Pif1 helicase, which translocates in the opposite (5’ to 3’) direction ^38^.

### Rep bypasses short dsDNA and DNA-RNA hybrid barriers

We next challenged Rep with defined nucleic acid barriers that it may encounter during its function. To generate these barriers, we annealed short, fluorescently labeled DNA or RNA oligonucleotides to the single-stranded λDNA (Figure 3A).

**Figure 3:**
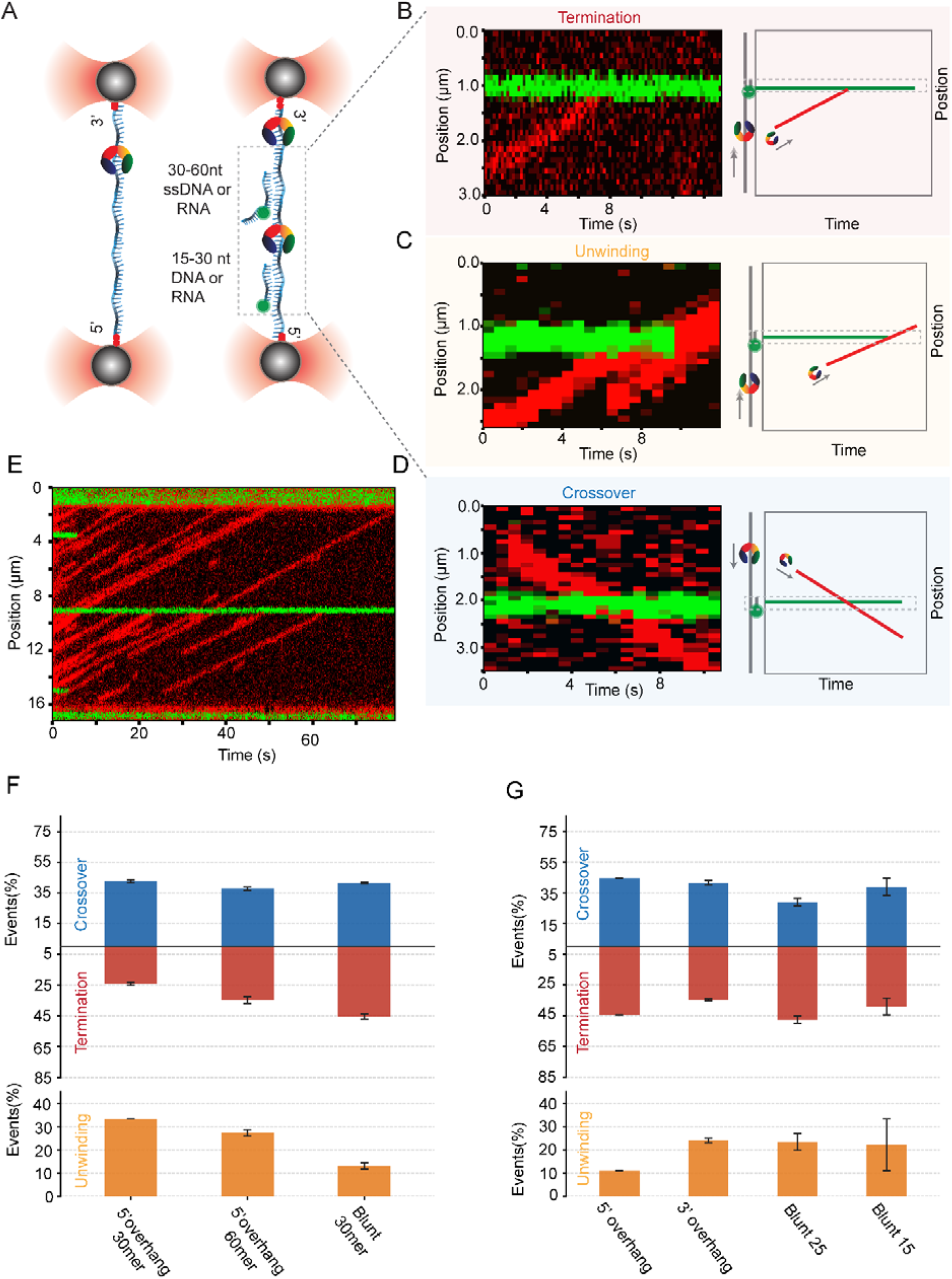
Rep can bypass, unwind, and terminate at DNA barriers. **A)** Schematic of the barrier assay. **B)** Example event of termination at the DNA barrier. **C)** Example event of unwinding at the DNA barrier. **D)** Example event of crossover at the DNA barrier. **E)** Example kymograph from a typical assay in which multiple events can be tracked simultaneously. **F)** DNA roadblock statistics. 24-mer 5^′^ overhang + 30-bp dsDNA (N = 33), 24-mer 5^′^ overhang + 60-bp dsDNA (N = 29), and blunt 30-bp duplex with no overhang (N = 53). Error bars were generated by bootstrapping. **G)** RNA roadblock statistics. 24-mer 5^′^ overhang + 30-bp hybrid (N = 27), 24-mer 3^′^ overhang + 30-bp hybrid (N = 58), blunt 25-mer hybrid (N = 21), and blunt 15-mer hybrid (N = 8). Error bars were generated by bootstrapping.

When a translocating Rep encountered a DNA-DNA or DNA-RNA duplex, we observed three distinct classes of events. In *termination* events, Rep dissociated upon reaching the duplex (Figure 3B). In *unwinding* events, probe fluorescence disappeared as Rep moved past the site, likely because of duplex unwinding (Figure 3C). In *bypass* events, Rep continued past the probe position while probe fluorescence remained detectable, indicating that Rep bypassed the short duplex without fully separating it from single-stranded λDNA (Figure 3D). Our confocal readout alone cannot distinguish passage over an intact duplex from transient unwinding followed by rapid reannealing behind the helicase; we return to this ambiguity below, where movement on long dsDNA resolves it. Four barrier sites were placed on one λDNA molecule to increase the frequency of observed encounters (Figure 3E).

For a 30⍰ bp DNA duplex with a 24-nt 5^′^ ⍰ overhang, apparent bypass was the most common (42 %), followed by unwinding (33 %) and termination (24 %) (Figure 3F). With a 60-bp duplex, the termination fraction increased and the putative unwinding fraction decreased (Figure 3F). Removing the 5’ overhang from the 30-bp duplex barrier increased termination and decreased unwinding, while keeping bypass at around 42 % (Figure 3F). In all three cases, a third or more of encounters resulted in apparent bypass while the relative frequencies of the other outcomes depended on barrier geometry.

These behaviors contrast with those of the closely related UvrD helicase examined using a comparable single-molecule approach. A translocating UvrD monomer is blocked upon reaching an ssDNA/dsDNA junction, where it stalls and then either dissociates or is joined, after a characteristic delay, by a second monomer that activates unwinding; stalled monomers never initiate unwinding on their own, and bypass of the duplex was not observed^26^. When two UvrD molecules reach the junction already associated, however, they unwind without a pause^26^. Rep differs in two respects: it frequently bypassed short duplex barriers, an outcome not observed for UvrD under comparable conditions, and in the subset of events that led to unwinding, unwinding began immediately rather than after a recruitment-limited delay. The absence of such a delay is consistent with an activating Rep assembly that is already present when the barrier is encountered. We note, however, that our measurements do not establish the oligomeric state of Rep responsible for either bypass or unwinding and determining the stoichiometry of the species underlying each behavior will be an important next step.

DNA-RNA hybrid barriers with a 3’ overhang, a 5’ overhang, or no overhang also produced all three operational outcomes, with apparent bypass in a third or more encounters for the conditions tested (Figure 3G). Across DNA-DNA and DNA-RNA substrates, continued Rep movement with persistent probe fluorescence was therefore reproducible.

### Rep translocates processively on double-stranded DNA

To ask whether Rep “bypass” behavior in Figure 3 might reflect bona fide translocation along dsDNA rather than unwinding, we engineered a construct containing an 69⍰nt ssDNA loading site followed by 48.5 kbp of λ⍰dsDNA, where the ssDNA represents a 3’ overhang (Figure 4A-B). Because Rep loads only onto ssDNA and cannot directly load onto duplex DNA (Figure 4C), this scheme ensures that any movement on dsDNA must arise from translocation initiated at the ssDNA overhang (Figure 4D). We observed that Rep consistently loaded adjacent to the bead (within the ssDNA region) and then translocated directionally along the dsDNA away from the loading site (Figure 4D–E). Translocation on dsDNA was highly continuous, exhibited no significant pauses, and average distance traveled was 12.7 kbp (n=54) (Figure 4F,4I).

**Figure 4:**
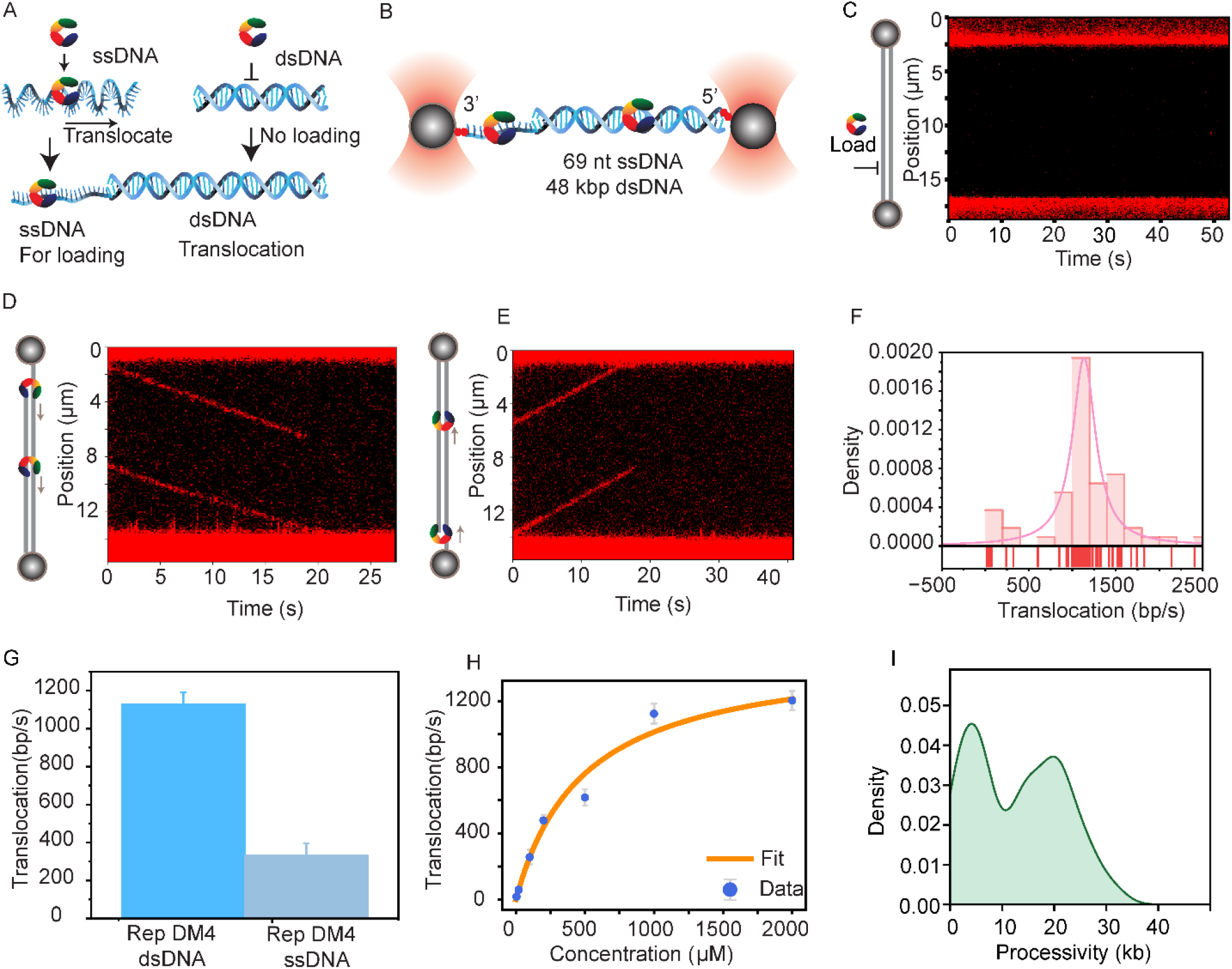
Rep translocates on dsDNA. **A)** Schematic of the dsDNA-translocation construct. An ssDNA segment is conjugated to λ-DNA to generate an ssDNA–dsDNA hybrid substrate.B)Schematic of assembled construct used for the experiment.C) Control experiment where Rep is loaded to only dsDNA construct where no terminal ssDNA is present. **D)** Representative kymograph of Rep translocating on dsDNA when the tether is held in the opposite orientation. **E)** Representative kymograph of Rep translocating on dsDNA when the tether is held in one orientation.F) Translocation rate of Rep on dsDNA (N = 54). **G)** Comparison of Rep translocation rates on ssDNA (N = 69) and dsDNA (N = 54). **H)** ATP titration for Rep translocation on dsDNA. ATP concentrations used: 20□µM (N = 30), 100□µM (N = 26), 200□µM (N = 18), 500□µM (N = 25), 1000□µM (N = 52), and 2000□µM (N = 26). I) Processivity of Rep on dsDNA for all the translocating traces evaluated under 1mM condition (n=54).

We observed a broad distribution of dsDNA translocation rates with a mean of 1100 bp/s at 1 mM ATP (Figure 4F-G), approximately 3.5 ⍰ fold higher than Rep’s translocation rate on ssDNA. An ATP titration yielded a K_m_ of 492.7 µM and a V_max_ of 1512 bp/s (Figure 4H). These observations resolve the ambiguity noted for the short-barrier bypass events above. Two independent arguments exclude unwinding masked by reannealing behind the helicase. First, kinetics: a process rate-limited by strand separation cannot exceed the unwinding rate (∼100 bp/s)^11^, yet dsDNA translocation (∼1100 bp/s) is roughly an order of magnitude faster and even exceeds ssDNA translocation (∼300 nt/s). Second, mechanics: unwinding at constant tension would shorten the tether and register as a force change, yet no tension change accompanies movement (Supplementary Figure 3). Together these indicate that Rep translocates along the duplex without separating the strands, and that the persistent-probe bypass events at short barriers most plausibly reflect the same duplex-tracking activity.

Overall, these results show that Rep overcomes duplex barriers through a combination of termination, unwinding, and most unexpectedly, directional and highly processive translocation on dsDNA. As with bypass and barrier unwinding, our data do not constrain the oligomeric state of the Rep species that translocates along duplex DNA; establishing the stoichiometry underlying each of these activities will be help place them within existing models of SF1A function.

### AlphaFold3 modeling suggests a strand-tracking mechanism on dsDNA

How might Rep translocate along duplex DNA? Rep, together with PcrA and UvrD, has long served as a structural prototype for SF1A helicases; the first helicase structures, in isolation and bound to DNA, were determined for Rep and PcrA. These studies established that a helicase monomer can translocate directionally along ssDNA, likely consuming one ATP per nucleotide, through an inchworm-like motion in which the 1A and 2A subdomains alternately close and open upon ATP binding and release. Whether this same machinery can operate on an intact duplex, however, has not been examined. As a first step toward a structural hypothesis, rather than a test of function, we used AlphaFold3 (AF3) to model Rep bound to a 30-bp duplex in the presence and absence of ATP (Figure 5).

**Figure 5:**
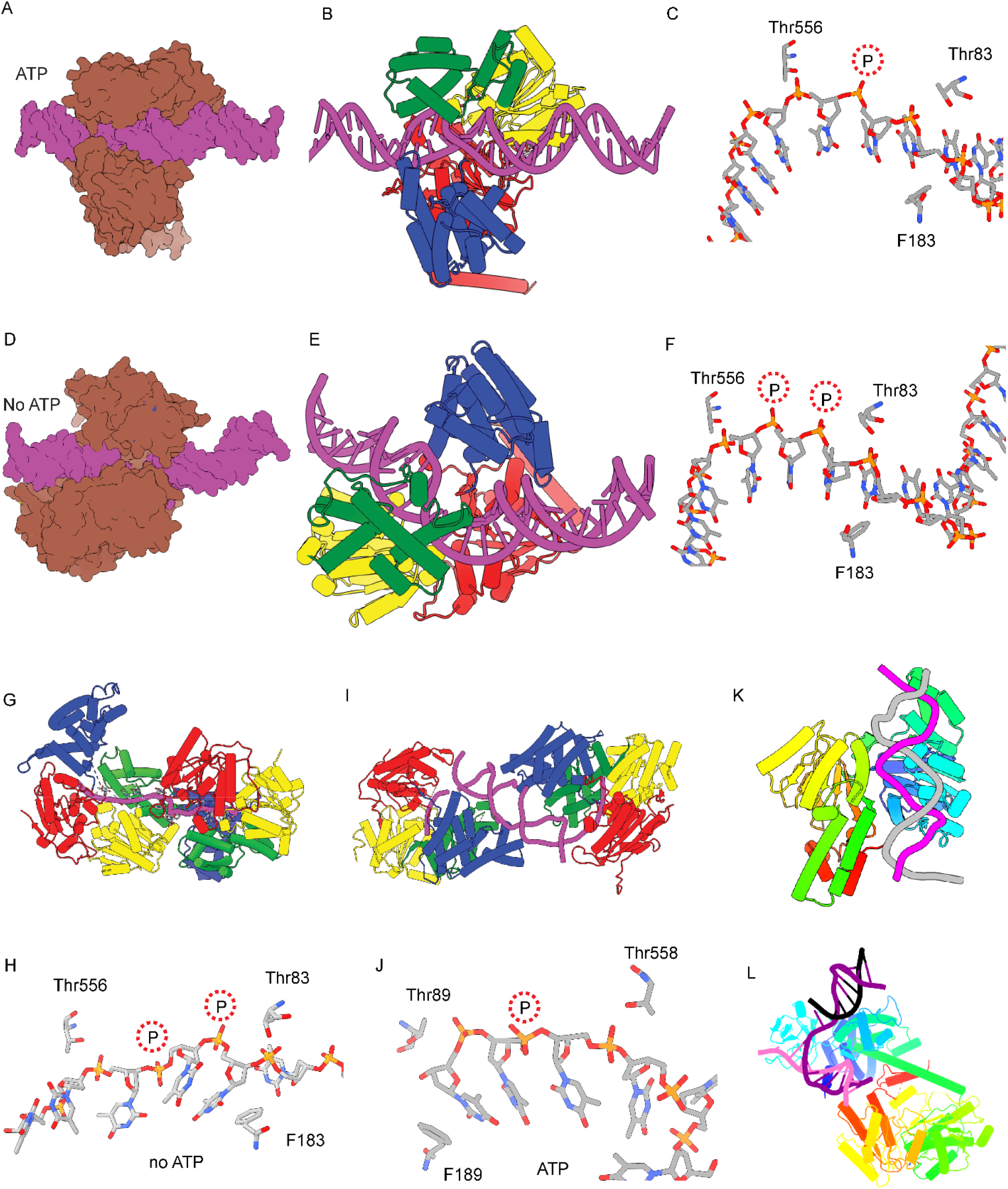
Computational modeling of Rep-dsDNA binding. **A–F)** AlphaFold3-predicted structures of Rep bound to a 30-bp dsDNA substrate. **A)** Surface projection of Rep (grey) bound to dsDNA (pink). **B)** Domain map of Rep, colored as in the tube representation, with dsDNA shown in pink. **C)** Zoomed-in view of the active site from B, highlighting Rep–dsDNA interactions. Protein residues are labeled by name; backbone phosphates are labeled as P. Panels A–C were predicted in the presence of ATP. **D–F)** AF3-predicted structures of Rep bound to dsDNA in the absence of ATP. **D)** Surface projection of Rep (grey) bound to dsDNA (pink). **E)** Domain representation of Rep (tubes) bound to dsDNA (tubes, pink). **F)** Zoomed-in view of the active site showing Rep–dsDNA interactions in the absence of ATP. **G)** Published structure of Rep bound to ssDNA, shown with the same labeling scheme (PDB: 1UAA). **H)** Zoomed-in view of the active site from G, showing Rep–ssDNA interactions during translocation. This structure was solved without ATP. **I)** Structure of UvrD, a Rep-like SF1 helicase, bound to forked DNA in the presence of an ATP analog (PDB: 2IS2), shown with the same schematic. **J)** Zoomed-in view of the active site from I, showing UvrD–ssDNA interactions during translocation. **K)** SF2 family helicase Rad54, which translocates on dsDNA, shown bound to dsDNA (PDB: 1Z63). DNA is colored by strand; protein is colored by default ChimeraX chain colors. **L)** Another representative dsDNA-binding SF2 helicase, RecG, shown bound to three-way DNA (PDB: 1GM5).

In both nucleotide states, AF3 placed dsDNA within Rep’s central groove, with the 1A and 2A motor subdomains engaging one strand while the complementary strand was accommodated without separation (Figure 5A-B), a geometry not represented in the ssDNA-bound structures on which such tools are trained. The motor subdomains reproduced the strand-specific contacts characteristic of ssDNA tracking: in the absence of ATP, Thr83 (1A) and Thr556 (2A) contacted backbone phosphates on the same strand three nucleotides apart, with Phe183 (1B) making some packing interaction in the major groove (Figure 5C). In the ATP ⍰bound model these contacts were retained but the two threonines moved to two nucleotides apart (Figure 5D–F). This ATP-dependent 3 nt to 2 nt change mirrors the threonine-spacing shift proposed to underlie one-nucleotide-per-ATP stepping in SF1 (Rep, UvrD) and SF2 (NS3, Vasa) helicases on single strands^39^.Moreover, under both nucleotide conditions, 2B domain makes additional contacts with the duplex DNA specifically Lys 493 contacting the major groove (Figure 5 and Supplementary Figure 5) in addition to the strand-tracking contacts. This 2B-mediated contact would likely help Rep stabilize duplex engagement while maintaining single-strand tracking.

To assess whether these interactions mirror those used during ssDNA translocation, we compared the models to the published Rep–ssDNA structure^22^ (Figure 5G). The same residues (Thr556, Thr83, and Phe183) engaged ssDNA with nearly identical spacing and orientation (Figure 5H), and UvrD structures showed the corresponding ATP-dependent changes in threonine-phosphate contacts (Figure 5I–J).

Finally, we compared these features to dsDNA⍰ bound structures of SF2 translocases. Rad54 tracks only one strand of the duplex despite binding both strands sterically (Figure 5K), and RecG similarly interacts primarily with one strand of the duplex arms in three ⍰ way junctions (Figure 5L). These parallels across SF1A and SF2 families support a general principle: duplex translocating helicases often track a single strand while the partner strand is passively accommodated.

Two caveats apply to these models. First, because AF3 is trained on existing helicase–nucleic-acid structures, its recapitulation of the known ssDNA contact geometry, including the ATP-dependent threonine spacing, is consistent with but does not independently establish that mechanism on dsDNA. The more informative prediction is the steric accommodation of an intact duplex alongside single-strand tracking, a geometry absent from the training structures. Second, these static models cannot address the dynamics, sufficiency, or processivity of translocation. We therefore interpret these models as a structural plausibility test rather than mechanistic validation, a hypothesis to be tested by future experimental structures.

## Discussion

For three decades, SF1A helicases have been understood as enzymes that track single-stranded DNA and unwind the duplexes they meet rather than travel along them. Using combined optical-trap and confocal fluorescence imaging of single Rep molecules on long DNA, we find that Rep instead transitions onto intact duplex DNA and translocates along it processively, and we use AlphaFold3 modeling to propose how such movement might occur.

Earlier stopped-flow and single-molecule measurements established Rep’s ssDNA translocation rate on short substrates but could not follow long-range movement. We reproduce that rate and show it is sustained continuously over kilobases, indicating that ∼300 nt/s is the intrinsic translocation rate and that the much slower unwinding reflects the energetic cost of strand separation.

A striking outcome is that a translocating Rep frequently bypasses a short dsDNA or DNA–RNA barrier, leaving the duplex intact behind it, in a third or more of encounters across all barrier types. Because Rep also translocates processively along long dsDNA, this bypass most likely reflects genuine movement along the duplex rather than the enzyme jumping over it. Such an activity would suit lagging-strand replication, where Okazaki fragments create frequent gaps and ssDNA–dsDNA junctions: rather than unwinding each intervening duplex, Rep could move efficiently between successive ssDNA regions while resolving secondary structures^44^ and redistributing bound SSBs, supporting rapid and faithful replication.

More broadly, processive dsDNA translocation may supply an activity that in vivo studies have implied but could not visualize: the clearing of nucleoprotein barriers ahead of the replication fork. Rep colocalizes with the fork, associates with DnaB in multiple copies, promotes replication restart, and helps replisomes traverse transcription units by removing stalled elongation complexes and other protein roadblocks^17–20,45-50^. Whether Rep reaches these obstacles by recruitment or from an adjacent ssDNA gap, movement along duplex DNA offers a mechanism for the barrier clearing these models require; consistent with this, Rep repositions SSB and RPA as it translocates on ssDNA, though we did not directly test protein displacement on the duplex. This assignment fits the contrasting cellular roles of Rep and UvrD. Rep is uniquely coupled to the replisome through DnaB and is the only accessory helicase whose loss slows chromosomal duplication^51^, whereas UvrD acts largely in post-arrest processing and repair^52^. That division is mirrored at a duplex: a UvrD monomer stalls at an ssDNA–dsDNA junction and does not bypass it^5,40^, whereas Rep both bypasses short barriers and translocates processively along long ones. Duplex-traversal capacity therefore tracks with the enzyme that acts as a dedicated, replisome-associated barrier-clearing motor.

dsDNA translocase activity is increasingly recognized across diverse DNA transactions, but until now within the SF2 and hexameric motors. In eukaryotes, many SF2 helicases act predominantly as dsDNA translocases, including Rad54 and the fork-reversal factors SMARCAL1, ZRANB3, and HLTF^7,9,53^; In bacteria, in bacteria, SF2 enzymes such as Mitomycin repair factor A (MrfA) and RecG translocate on dsDNA during repair^7,54-55^. To our knowledge, Rep is the first SF1 helicase shown to translocate processively along duplex DNA, extending this capability to the SF1 family. The same principle operates on RNA: the SF2 RIG-I-family helicases RIG-I, MDA5, and LGP2 translocate along double-stranded RNA^56-58^, indicating that duplex translocation is a mechanism shared across both nucleic acids and multiple helicase superfamilies.

Beyond its activity on duplex DNA, Rep also drives the directional movement of ssDNA-binding proteins along ssDNA. Related behavior has been reported for other motors: the structural homolog yeast Pif1 redistributes human RPA in its direction of translocation^38^, the E. coli helicase YoaA relocates SSB^59^, and PriA has been proposed to redistribute SSB on the lagging strand^17^. Our results add Rep to this group and, because Rep redistributes bacterial SSB, archaeal SSB, and eukaryotic RPA, indicate a robust, largely kinetic mechanism in which directional translocation biases the diffusion of the bound protein rather than relying on specific protein–protein contacts.

Such redistribution could serve several roles. By clearing SSB from the lagging strand, Rep could expose ssDNA for loading of replication-restart machinery; more broadly, a translocase that displaces bound proteins parallels the anti-recombinase activities of UvrD and Srs2, which dismantle RecA or RPA filaments during recombination^60–63^. We previously showed that repetitive shuttling of Rep on a stalled-fork-like substrate can delay RecA filament formation on ssDNA^15^, consistent with a role in keeping ssDNA tracts clear of bound proteins.

How might Rep move along an intact duplex? Our AF3 models suggest that its canonical stepping mechanism can operate with the duplex left intact: the motor tracks a single strand through the same one-nucleotide-per-ATP inchworm cycle it uses on ssDNA, while the complementary strand is passively accommodated rather than separated. Initial engagement likely occurs through the 2B subdomain, which our models place in contact with the duplex and which prior work shows mediates dsDNA contacts in the related helicases UvrD^5^ and PcrA^64^, consistent with Rep engaging dsDNA more efficiently in the presence of AMPPNP^41^. This fits the emerging view of 2B as a conformational switch that gates SF1A activity: a large reorientation of 2B accompanies activation for unwinding in an SF1A dimer^65^, the same 2B interface governs dimerization and processivity across UvrD, Rep, and PcrA^13^, and constraining the 2B subdomain of a monomeric Rep by intramolecular crosslinking converts it into a processive superhelicase^11^. Together these indicate that the positioning and mobility of 2B, rather than any single fixed contact, selects a given activity, and that duplex tracking without unwinding likely represents an additional setting of this switch.

Notably, translocation on dsDNA is approximately 3.5-fold faster than on ssDNA, suggesting a more slippery, mobility-optimized mode of movement mediated by weaker contacts with the backbone. Consistent with a role for the 2B subdomain in setting speed, the RepΔ2B variant translocates ∼2-fold faster than wild-type Rep on ssDNA^14^, indicating that 2B engagement can tune the rate of movement. This dsDNA translocation rate is comparable to the replication rate in E. coli, so if Rep clears nucleoprotein obstacles ahead of the fork by translocating on the duplex, its speed would be physiologically appropriate.

Our demonstration that a canonical single-stranded DNA translocase can traverse an intact duplex broadens the mechanistic repertoire attributed to SF1 helicases and invites re-examination of how these motors operate within the crowded environment of replication and repair. Whether this capability extends to other SF1A helicases remains open and must be tested enzyme by enzyme: the closely related UvrD does not bypass an ssDNA–dsDNA junction^40^, showing that duplex-translocation competence is not a given even among near relatives.

## Author Contributions

T.H, S.P, and O.Y conceived the project. S.P, A.M.N, O.Y performed the optical trap experiments. S.P prepared and labeled the Rep protein and DNA constructs with the inputs from O.Y. F.R and J.M.B prepared SsoSSB. S.P analyzed the data, prepared the draft and wrote the ms with T.H. All authors helped with writing, editing and finalizing the ms.

## Acknowledgements

We would like to thank Tim Lohman for providing SSB labeled protein as a gift and Edwin Antony for providing RPA as a gift. We would like to thank the team at LUMICKS for their technical help, especially Suresh Ramakoti. Authors would like to thank all the members of Ha and Myong labs along with Fire, Aether, morning coffee, and Friday Happy hour subgroups for their helpful discussion and suggestions for this project.

