## Supplementary files for "A superfamily 1 helicase translocates on double-stranded DNA"

Methods

Protein Purification and labeling

Rep was prepared the same as previously described. pET28a(+) with rep(C18L/C43S/C167V/C612A/S400C) was transformed into E. coli B21(DE3) (Sigma-Aldrich, CMC0014). A single colony was picked and grown in TB at 37°C overnight, followed by 30°C overnight. When OD reached the range between 0.3 and 0.4, the cells were moved to an 18°C incubator. When OD reaches 0.6 to 0.8, the cells were induced expression with 0.5 mM IPTG and continue growth overnight. The cells were harvested by centrifugation for 15 min at 10,700 × g and 4°C. The pellet was resuspended in 40 ml of the lysis buffer (50 mM Tris-HCl pH 7.5, 5 mM Imidazole, 200 mM NaCl, 20% (w/v) sucrose, 15% (v/v) glycerol, 17.5 μg/ml PMSF, and 0.2 mg/ml Lysozyme) and sonicate to lyse the cells. The lysed cell mix was centrifuged at 30,000 × g at 4°C for 30-60 min and collect the supernatant. The supernatant was stir-mixed with pre-equilibrated Ni-NTA resin for 1.5 h at 4°C. Ni-NTA purification was performed by washing the protein-bound resin with buffer A (50 mM Tris-HCl pH 7.5, 5 mM Imidazole, 150 mM NaCl, 25% (v/v) glycerol), followed by buffer A1M (50 mM Tris-HCl pH 7.5, 5 mM Imidazole, 1 M NaCl, 25% (v/v) glycerol) to remove any DNA residue, and final washed the protein-bound resin with buffer A, then eluted the Rep variant with imidazole buffer (50 mM Tris-HCl pH 7.5, 205 mM Imidazole, 150 mM NaCl, 25% (v/v) glycerol). The excess Imidazole was removed by an overnight dialysis and stored in Rep storage buffer (50% glycerol, 600 mM NaCl, 50 mM Tris-HCl pH 7.5) at −80°C. Proteins were labeled using Cy3 and Cy5-malemide. Briefly, dyes were mixed with protein in 10X excess and incubated overnight in 4C. Then excess dyes were removed through dialysis using 3.5KDa filter dialysis bags.

SsoSSB purification and labeling

Untagged *Sulfolobus solfataricus* SSB-A114C was expressed from pRSF-SsoSSB-A114C in BL21-AI cells grown in 2×YT containing 50 µg/mL kanamycin at 37°C to an OD₆₀₀ of approximately 0.8. Expression was induced with 2 g/L L-arabinose and 0.5 mM IPTG, and the culture was incubated for 6 hours at 30°C. Cells were harvested and resuspended in lysis buffer containing 50 mM HEPES (pH 7.5), 200 mM NaCl, 15% glycerol, 2 mM DTT, and protease inhibitors. Following treatment with 2 mg/mL lysozyme for 30 min on ice, cells were disrupted by sonication and the lysate was clarified by centrifugation at 18,000 rpm for 30 min at 4°C. The soluble fraction was heated at 70°C for 20 min and centrifuged again at 20,000 rpm for 20 min at 4°C to remove heat precipitated proteins. The heat-soluble fraction was diluted to approximately 100 mM NaCl and applied to a HiTrap Heparin HP column equilibrated in 50 mM HEPES (pH 7.5), 100 mM NaCl, 15% glycerol, and 1 mM TCEP. Bound proteins were eluted with a linear gradient from 100 mM to 1 M NaCl, and fractions from the elution peak containing SsoSSB-A114C were pooled and concentrated. The protein was further purified on a HiLoad 16/600 Superdex 75 pg column equilibrated in 50 mM HEPES (pH 7.5), 500 mM KCl, 15% glycerol, and 1 mM TCEP. Purified SsoSSB-A114C was labeled by incubation with a 10-fold molar excess of Alexa647N-maleimide, with the reaction protected from light. Unreacted dye was removed using a PD-10 desalting column, and the labeled protein was further purified by a second Superdex 75 size-exclusion chromatography step to remove residual free dye. Fractions containing Alexa647N-labeled SsoSSB-A114C were pooled, concentrated, divided into single-use aliquots, snap-frozen in liquid nitrogen, and stored at −80°C

Ensemble FRET Assays

The ensemble FRET unwinding assay was performed by assembling reactions in a cuvette to final concentrations of 10 mM Tris, 10 mM MgCl₂, 5–15 mM NaCl (accounting for salt carried with the helicase), 1 mM ATP, 1% BSA, 5 nM DNA substrate, and 10 nM helicase. A 2.5x reaction buffer (25 mM Tris pH 8.0, 25% glycerol, 25 mM MgCl₂) was mixed with NaCl, BSA, and ddH₂O, excluding DNA, helicase, and ATP, and loaded into the fluorimeter cuvette. Using the Cary Eclipse in Kinetics mode, the cuvette was equilibrated until the fluorescence signal stabilized. DNA was added and mixed rapidly, followed by equilibration, then helicase (diluted to 1 µM in cold 1 M NaCl for solubility), which produced a small PIFE‑associated signal change. After the signal restabilized, ATP was added to initiate unwinding.

dsDNA Lambda (λ) and ssDNA-dsDNA construct preparation

λ DNA (48.5 kb with 12-nucleotide 5’ overhangs) was purchased from NEB. A series of annealing and ligation steps were performed to generate a construct with biotin modifications on the same strand, as previously described^37,66^.

For ssDNA-dsDNA construct similar strategy was used. Same lambda DNA was used with 12-nucleotide 5’ overhangs. In brief, four oligonucleotides Oligo 1, Oligo 2’, Oligo-Rep and Oligo 3 were synthesized by Integrated DNA Technologies Inc. (IDT). Oligo 1, Oligo-Rep and Oligo 3 were then phosphorylated to prepare them for ligation. Phosphorylation was performed using T4 PNK (NEB) at 37°C for 1 h. Next, Oligo 1 and Oligo 3 were annealed to the 5’ overhangs by heating to 65°C, followed by slow cooling. Immediately after annealing, ligation was performed using T4 DNA ligase (NEB) at room temperature for 20 min, resulting in biotinylation of one 3’ end of the DNA. Subsequently, Oligo 2’ and Oligo-Rep were annealed by heating to 95°C and slowly cooling down to 4°C at a rate of 1°C reduction per 30 seconds. Resulted annealed product was then annealed to the previously ligated product by heating to 45°C, followed by slow cooling. Another ligation step was performed at room temperature for 20 min. To remove excess primers and proteins, drop dialysis was performed using 0.025 µm filters (Millipore Sigma). A membrane was carefully suspended over ~20–25 mL of dialysis buffer (10 mM Tris-HCl, pH 8.0, and 0.1 mM EDTA) in a Petri dish. A drop of DNA (100–150 µL) was placed onto the membrane using a large-orifice pipette tip, ensuring that the final volume remained within this range to prevent the membrane from dipping into the buffer. Dialysis was carried out at room temperature for at least 1.5 h. Afterward, the solution was carefully collected using large-orifice pipette tips. DNA was then stored in 4°C.

Single Molecule Optical Tweezers Assay

A LUMICKS C-trap, a commercial optical trap with combined confocal microscopy, was used for all single molecule measurements. The passivation and cleaning protocol were followed in exact same was as is explained in detail here^65^. Briefly, streptavidin-coated polystyrene beads (4.34 µm) were used to capture λ-DNA or junction λ-DNA. For ssDNA, λ-DNA was mechanically denatured to generate ssDNA and verified by fitting force-distance curves to the FJC model. Experiments were performed in 100mM NaCl 25mM Tris-HCL pH8.0,2 mM MgCl₂ and ATP, with an oxygen-scavenging system (dextrose, glucose oxidase, catalase, and Trolox) to enhance fluorophore stability. Imaging was conducted with 488, 561, and 638 nm lasers at 100 nm pixel size and 0.1-0.5 ms pixel exposure, maintaining 5 mW power at the objective.

**Data Analysis**

All single molecule data were analyzed using custom scripts based on Lumicks Pylake python package. Analysis scripts have been deposited in the [Github](https://github.com/spangeni/DnaB-project). Plots were generated using Python, Origin Pro 2024b version and illustrator.

Supplementary Figures:

Supplementary Figures:

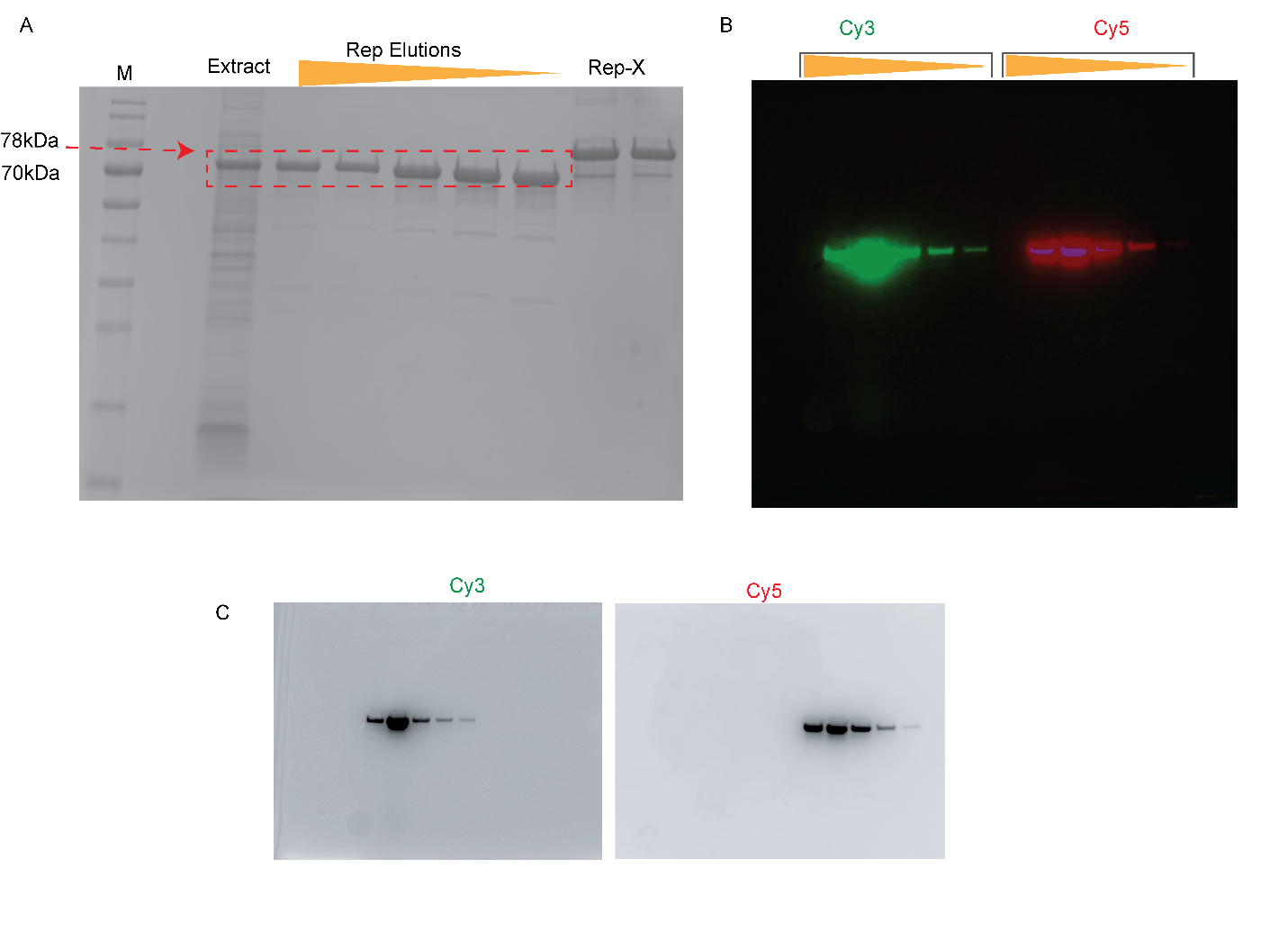

***Supplementary Figure 1. Purification, fluorescent labeling, and imaging of Rep helicase. A)*** *Rep was purified from E. coli expression strains and analyzed by SDS-PAGE to assess protein purity. B) Purified Rep was subsequently labeled with Cy3 and Cy5 fluorophores and characterized by column chromatography, with elution fractions analyzed to confirm successful labeling. C) Representative fluorescence images of labeled Rep acquired in the individual Cy3 and Cy5 channels demonstrate fluorescent signal detection from the labeled protein.*

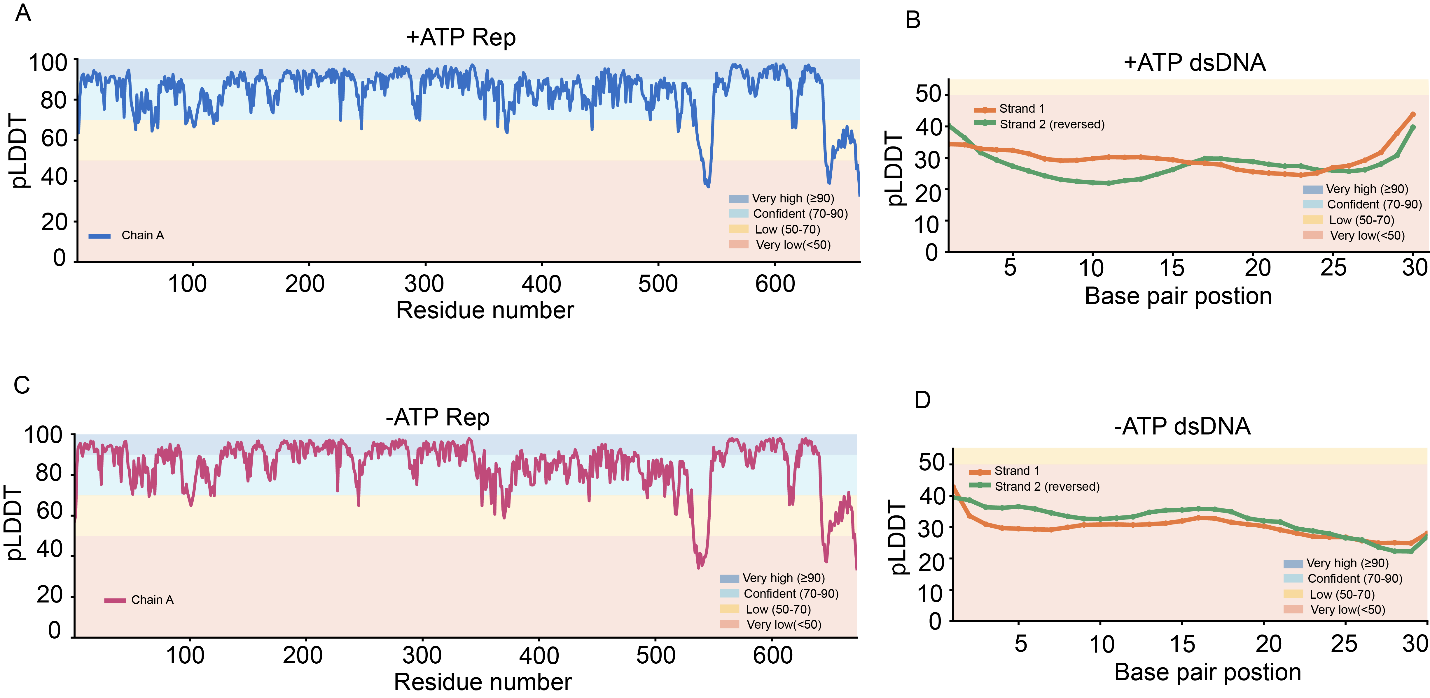

***Supplementary Figure 2. AlphaFold3 predictions of Rep bound to dsDNA in the presence and absence of ATP.A)*** *pLDDT confidence scores are shown for AlphaFold3 (AF3) models of Rep in complex with dsDNA. In the presence of ATP and Mg²⁺, Rep is predicted with uniformly high confidence, consistent with the availability of experimentally determined structures and the well-folded nature of the protein. B) In contrast, the dsDNA chains exhibit lower pLDDT values, reflecting the reduced ability of AlphaFold-style models to accurately predict DNA conformations in isolation rather than indicating structural disorder. C-D) Similar trends are observed in the absence of ATP. Rep remains highly confident throughout the protein structure (****C****), while the DNA chains are predicted with lower confidence scores, as observed for the ATP-bound model (****D****).*

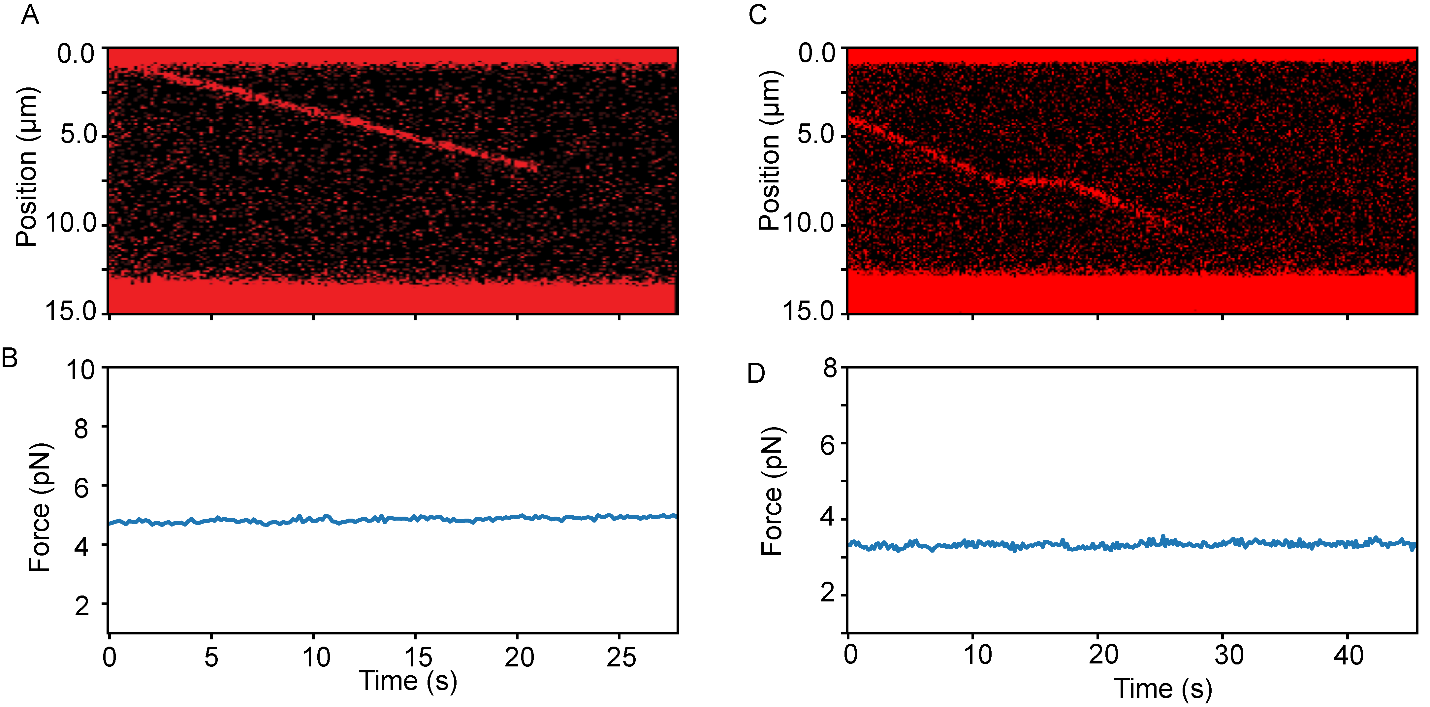

***Supplementary Figure 3: Rep dsDNA translocation traces combined with force****. A-B) A translocation trace of Rep at 5pN. No visible force changes observed suggesting no unwinding. C-D) A translocation trace of Rep where a transient pause was observed at 3pN. No visible force changes observed suggesting no unwinding.*

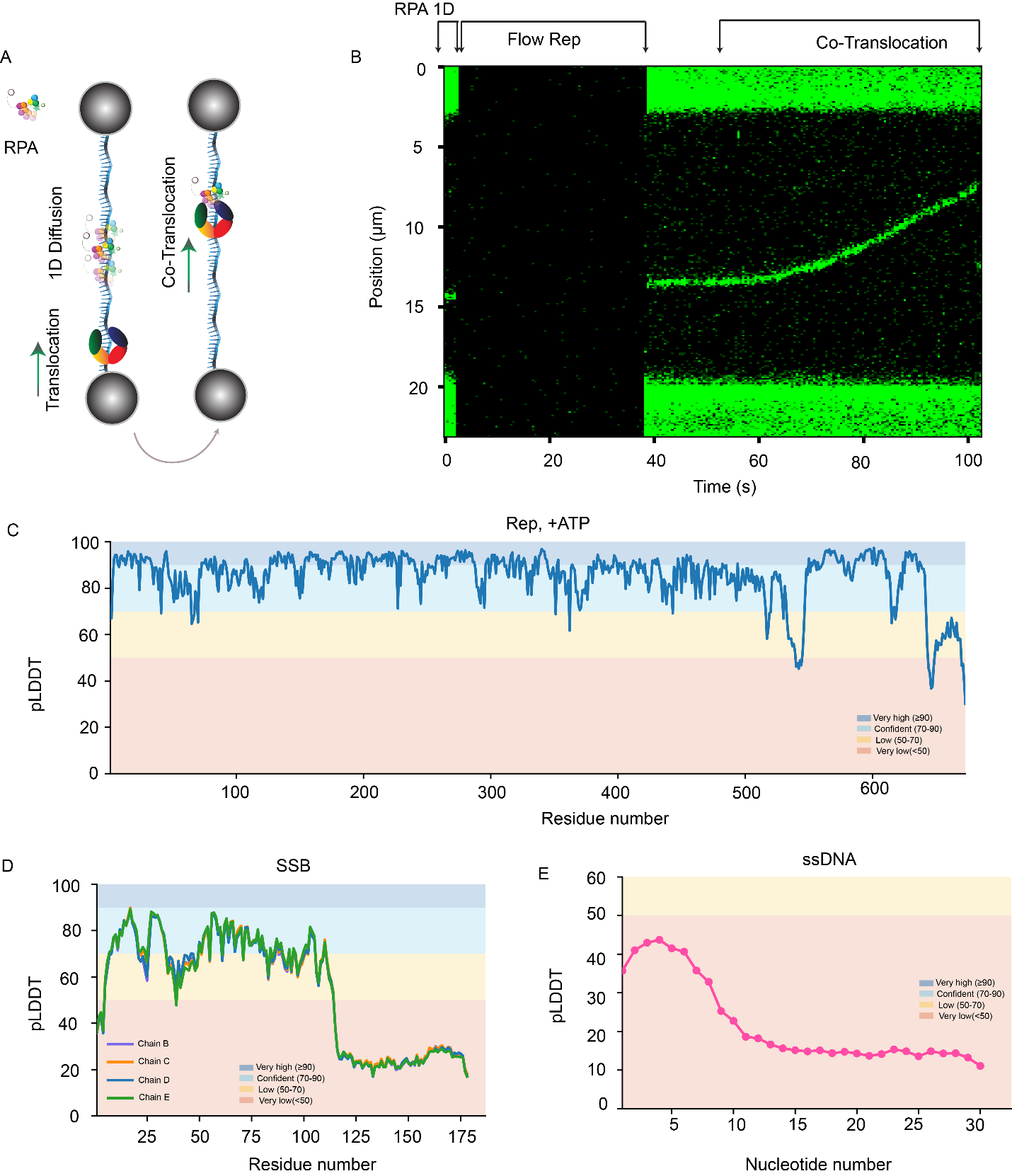

***Supplementary Figure 4. Rep interaction with RPA on ssDNA and AlphaFold3 predictions of the Rep-RPA-ssDNA complex.*** *A) A schematic of the Rep-RPA encounter assay is shown, in which RPA diffuses along ssDNA while Rep translocates directionally on the same substrate.B) A representative kymograph illustrating a Rep-RPA encounter event. In this assay, RPA is fluorescently labeled, whereas Rep is unlabeled.C-E) AlphaFold3 (AF3) pLDDT confidence scores for the Rep-RPA-ssDNA complex. C) Rep is predicted with high confidence throughout most of the protein structure .D) RPA, which functions as a heterotrimeric complex, is also predicted with high confidence across its structured domains, while flexible and unstructured regions display lower confidence scores as expected .E) The ssDNA chain exhibits overall lower pLDDT value, reflecting the inherent difficulty of predicting flexible single-stranded nucleic acid conformations rather than indicating a lack of biological relevance.*

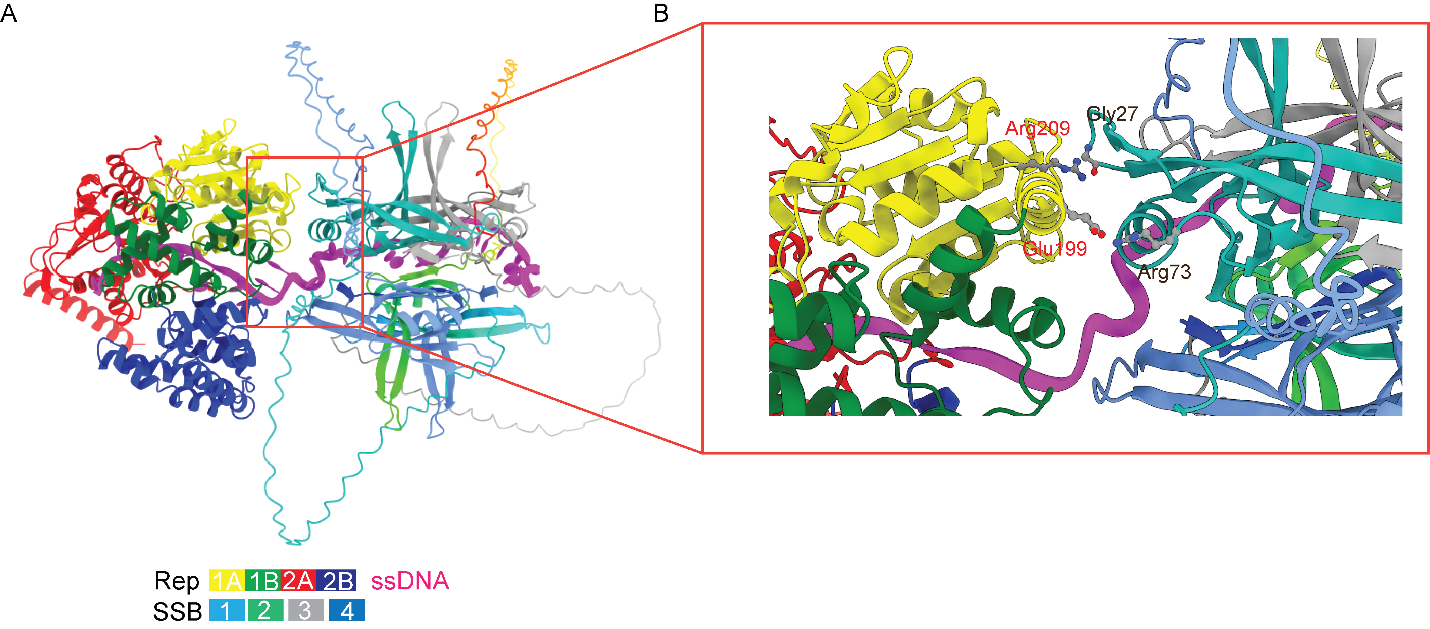

***Supplementary Figure 5****:* ***Rep-SSB-ssDNA-ATP structure prediction with AF3****. A) Assembled structure as per AF3 model prediction as in Figure2. B) Zoomed in version of transient interaction residues spanning the 1A domain of Rep and monomer of SSB.*

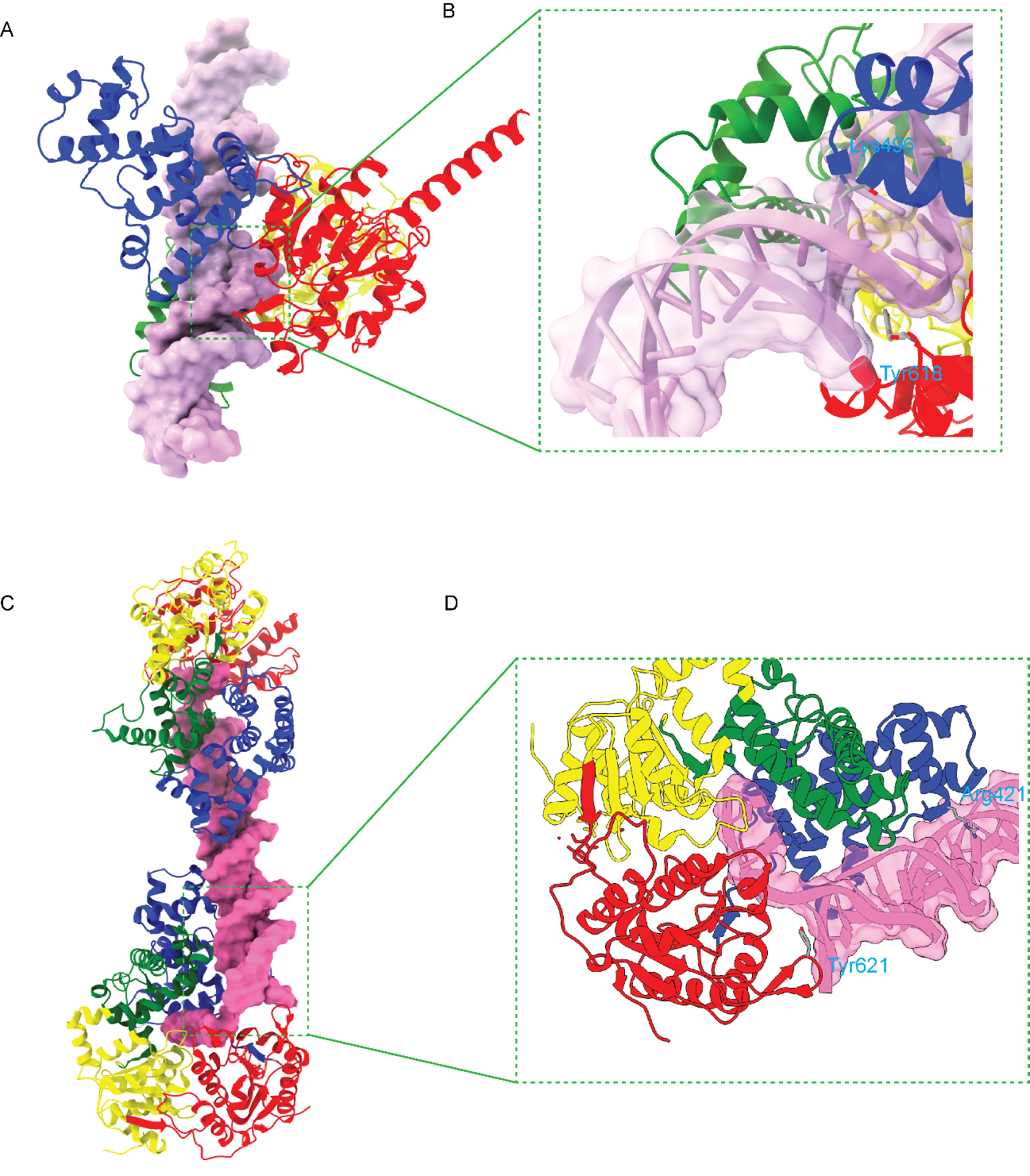

***Supplementary Figure 6. Comparison of dsDNA contact sites in the SF1A helicases Rep and UvrD.*** *A-B) AlphaFold3 (AF3)-predicted structures of Rep bound to dsDNA. The overall Rep-dsDNA complex is presented in* ***(A)****, while a magnified view of the DNA-contacting region is shown in* ***(B)****. The zoomed view highlights interactions between dsDNA and residues Lys496 in the 2B domain and Tyr618 in the 2A domain of Rep.C-D) An analogous DNA-bound structure of the closely related SF1A helicase UvrD (PDB: 2IS2). The overall UvrD-junction DNA complex is presented in* ***(C)****, with a magnified view of the DNA-contacting interface shown in* ***(D)****. Interactions are observed in UvrD, where Arg421 in the 2B domain and Tyr261 in the 2A domain contact the DNA substrate.*

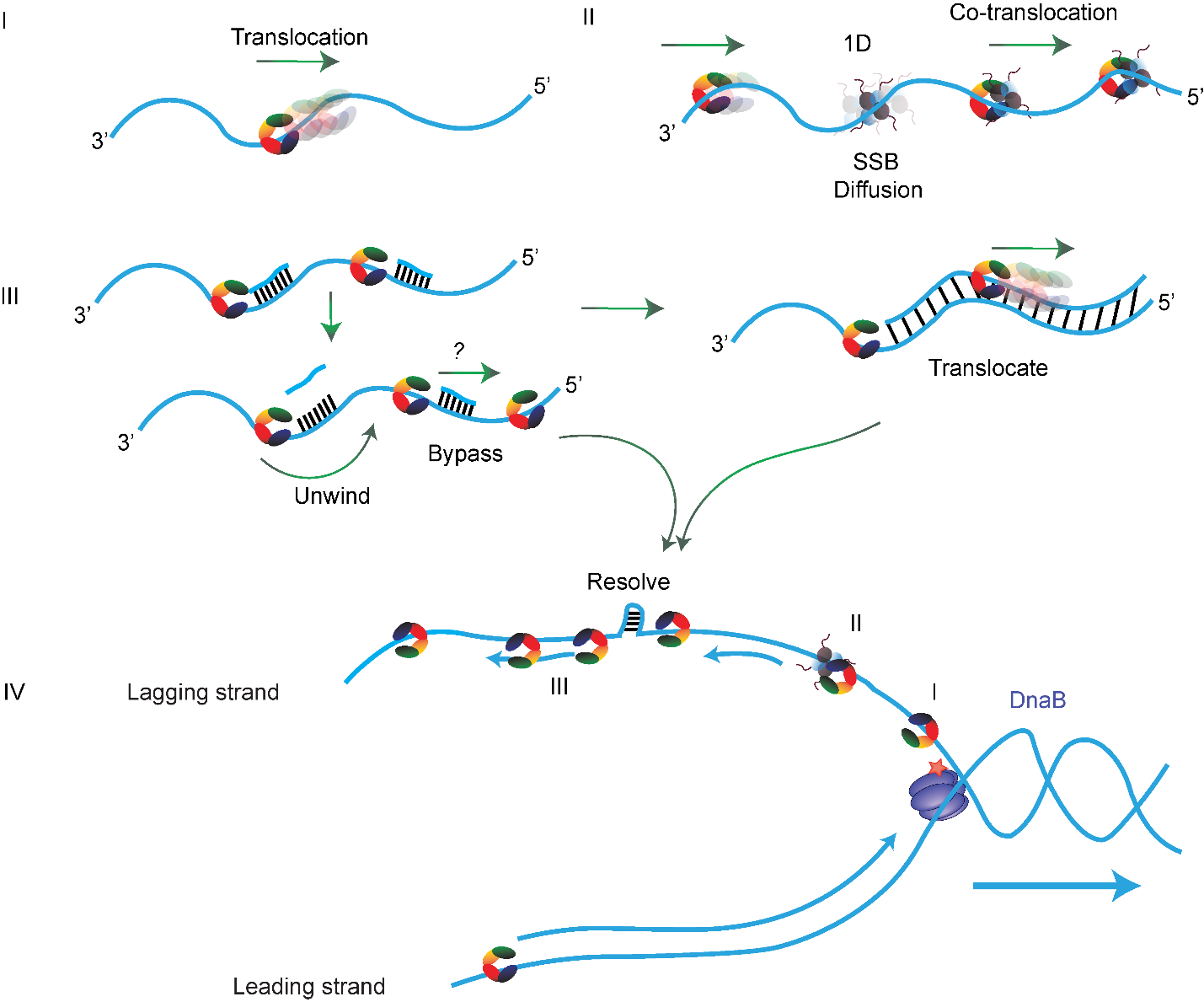

***Supplementary Figure 7. Proposed model for Rep activity during DNA replication and repair.*** *I)Rep loads onto ssDNA and translocates directionally. II) During translocation, Rep can drive the movement of diffusively bound SSB proteins along ssDNA. III) When encountering short dsDNA barriers, Rep may either unwind or bypass them; in the presence of extended blunt-ended dsDNA, Rep may transition onto and translocate along dsDNA. IV) These activities may facilitate lagging-strand replication by redistributing SSB, resolving secondary structures, and overcoming obstacles such as short duplex regions and Okazaki fragments.*

Supplementary Table 1: Oligos used in the construct preparation and experiments

| Name | Sequence 5’-3’ |
| --- | --- |
| Oligo 1 | ggg cgg cga cct gga caa |
| Oligo-Rep | aggtcgccgcccttttttttttttttttttttttttttttttttttttttttttttttttttttttttttttttttttatgcgctagtac |
| Oligo 3 | ibiodt /t/ ibiodt/ t/ ibiodt/t ttt ttt aga gta ctg tac gat cta gca tca atc ttg tcc |
| Oilgo 2 | gta cta gcg cat ttt ttt t/ibiodt/t /ibiodt/t/3bio/ |
| Cy3 probe | gcg gta gtc gcc /3cy3sp/ |
| Oligo 30_a | gat gtt ctg ctg gat atg cac ttt tcc ggg ggc gac tac cgc tgc ctg ggt gca |
| Oligo 30_b | gca gcc aca gtc act cat tgt ccg gta cag tgc ctg ggt gca ggc gac tac cgc |
| Oligo 30_c | cgc cca tat cat cca gtg gtc gta gca gtc ggc gac tac cgc tgc ctg ggt gca |
| Oligo 30_d | caa tgt cgc ccc aag acc atc tct atg agc tgc ctg ggt gca ggc gac tac cgc |
| Oligo60_a | cag tac cca tta ttt tca tga tgt tct gct gga tat gca ctt ttc cgg gct gac gta cac ggc gac tac cgc tgc ctg ggt gca |
| Oligo60_b | gtt acg tct gca tgt gct atc tgc gcc cat atc atc cag tgg tcg tag cag tcg ttg atg ggc gac tac cgc tgc ctg ggt gca |
| Oligo60_c | ggt tcc tgc ata tga tga caa tgt cgc ccc aag acc atc tct atg agc tga aaa aga aac tgc ctg ggt gca ggc gac tac cgc |
| Oligo60_d | gga cag tca ggg cag cca cag tca ctc att gtc cgg tac agc cgt gag tgc cgg tgc aca tgc ctg ggt gca ggc gac tac cgc |
| RNA_3’ | /5cy3/rcrgrgrcrurgrgrcrurgrgrurgrararcrururcrcrgrarurargrurgrcrgrgrgrurgrururururururururururururururururururururururururururu |
| RNA_blunt_1 | rgrgrcrurgrgrcrurgrgrurgrararcrururcrcrgrarurargrurgrc/3cy3sp/ |
| Ensemble_top | tg gcg acg gca gcg agg cat gtt aat act tct cct atg act gct agt tga (amino-dt-cy3) t29 |
| Ensemble_bot | agt att aac atg cct cgc tgc cgt cgc ca |
| Blunt_30 | gatgttctgctggatatgcacttttccggg/3cy5sp/ |
| RNA_5’ | rururururururururururgrgrcrurgrgrcrurgrgrurgrararcrururcrcrgrarurargrurgrc/3cy3sp/ |
| RNA_blunt_2 | rgrgrcrurgrgrcrurgrgrurgrararcruru/3cy3sp/ |

**Alpha Fold structures:**

Rep:MRLNPGQQQAVEFVTGPLLVLAGAGSGKTRVITNKIAHLIRGSGYQARHIAAVTFTNKAAREMKERVGQTLGRKEARGLMISTFHTLGLDIIKREYAALGMKANFSLFDDTDQLALLKELTEGLIEDDKVLLQQLISTISNWKNDLKTPSQAAASAIGERDRIFAHVYGLYDAHLKACNVLDFDDLILLPTLLLQRNEEVRKRWQNKIRYLLVDEYQDTNTSQYELVKLLVGSRARFTVVGDDDQSIYSWRGARPQNLVLLSQDFPALKVIKLEQNYRSSGRILKAANILIANNPHVFEKRLFSELGYGAELKVLSANNEEHEAERVTGELIAHHFVNKTQYKDYAILYRGNHQSRVFEKFLMQNRIPYKISGGTSFFSRPEIKDLLAYLRVLTNPDDDCAFLRIVNTPKREIGPATLKKLGEWAMTRNKSMFTASFDMGLSQTLSGRGYEALTRFTHWLAEIQRLAEREPIAAVRDLIHGMDYESWLYETSPSPKAAEMRMKNVNQLFSWMTEMLEGSELDEPMTLTQVVTRFTLRDMMERGESEEELDQVQLMTLHASKGLEFPYVYMVGMEEGFLPHQSSIDEDNIDEERRLAYVGITRAQKELTFTLAKERRQYGELVRPEPSRFLLELPQDDLIWEQERKVVSAEERMQKGQSHLANLKAMMAAKRGK

DNA top strand: TTTTTTTTTTTTTTTTTTTTTTTTTTTTTT

DNAA complementary strand: AAAAAAAAAAAAAAAAAAAAAAAAAAAAAA

Ion: Magnesium (Copy (1)

Ligand: ATP (Copy1)

SSB:MASRGVNKVILVGNLGQDPEVRYMPNGGAVANITLATSESWRDKATGEMKEQTEWHRVVLFGKLAEVASEYLRKGSQVYIEGQLRTRKWTDQSGQDRYTTEVVVNVGGTMQMLGGRQGGGAPAGGNIGGGQPQGGWGQPQQPQGGNQFSGGAQSRPQQSAPAAPSNEPPMDFDDDIPF(Copy 4)

ssDNA: TTTTTTTTTTTTTTTTTTTTTTTTTTTTTT
